# Water use efficiency of *Cupressus gigantea* is higher in mountain slope habitats compared to riverside habitats

**DOI:** 10.1101/2024.09.25.615050

**Authors:** Yueliang Jiang, Li Luan, Youlu Zuo, Shengyun Liu, Qiaoling He, Jingyu He, Ruru Wu, Gaoming Jiang, Meizhen Liu

**Affiliations:** POWERCHINA Chengdu Engineering Corporation Limited, Chengdu 611130, China; State Key Laboratory of Vegetation and Environmental Change, Institute of Botany, the Chinese Academy of Sciences, Beijing 100093, China; College of Resources and Environment, University of the Chinese Academy of Sciences, Beijing 100049, China; Graduate School of Chinese Academy of Sciences, University of the Chinese Academy of Sciences, Beijing 100049, China

**Keywords:** *Cupressus gigantea*, Water use efficiency, Stable isotope, Water source

## Abstract

*Cupressus gigantea* is an endemic species of the Yarlung Zangbo River in Tibet, China, and is designated as a national key protected wild plant. It is only distributed in the narrow area along Yarlung Zangbo River from Jiacha County to Danniang village Milin City in various habitats such as the waterline along the Yarlung Zangbo River, terraces, and mountain slopes. In this study, we utilized stable isotope technology (δD, δ^18^O, and δ^13^C), combined with the IsoSource model, to quantitatively determined the sources of water absorption and the water use efficiency of *C. gigantea* in various habitats, including riverside area, terraces and mountain slopes. The results showed that the content of soil total nitrogen, soil nitrate nitrogen, soil ammonium nitrogen, and soil organic carbon are significantly higher on mountain slope than in the other two habitats, while the total carbon content, water content, and total nitrogen content of *C. gigantea* leaves are significantly lower on mountain slope than in the other two habitats (*P*<0.05). The water use efficiency (WUE) of *C. gigantea* in the mountain slope habitats is 107.8 μmol mol^-1^, which is significantly higher than that of other habitats (*P*<0.05), with the WUE of *C. gigantea* in the riverside habitat being the lowest at 35.7 μmol mol^-1^. *C. gigantea* distributed on the mountain slope and terrace uses soil water as the main water source. During the normal rainy season, the proportion of soil water utilization is the highest, accounting for more than 60%; in the dry season, the utilization of river water and groundwater increases, while soil water utilization decreases. *C. gigantea* distributed along the riverside mainly absorbs water from the Yarlung Zangbo River, about 63%. In August when the rainfall increases in the rainy season, the use of soil moisture increases, and in the dry season, river water becomes the main water source. This indicates that with the alternation of rainy and dry seasons, the water absorption sources and strategies of *C*.*gigantea* adjusted accordingly. The research results provide an important scientific basis for the protection and management measures of the *C. gigantea* population.

## INTRODUCTION

The sources of water absorbed by plants generally include soil water, river water, groundwater, atmospheric precipitation, and various other sources. The proportion of hydrogen and oxygen isotope ratios in water varies among these sources. Therefore, it is possible to analyze which sources of water are used by comparing and analyzing the isotope content of water in different water bodies and plants. This analysis can directly reflect the differences in the source of water absorbed by plants in response to environmental changes, as well as the changes in vegetation water use, adaptability to the natural environment, and ecological stress that plants experience (*Ma & Song et al., 2016; Geris et al., 2017*). The IsoSource model was used to calculate the relative utilization of multiple water sources by plants, as well as the contribution rate and range of these water sources to plant water (*Gregg et al., 2003; Phillips et al. 2003*). In the mountainous areas of California along the banks of perennial rivers, the early transpiration of trees during the growing season mainly comes from the soil water, while during most of the dry seasons, it comes from groundwater (*Smith et al. 1991*); Studies on the soil-water-atmosphere continuum indicate that the soil depth at which root water uptake occurs largely depends on seasonal climate fluctuations, as observed using stable isotopes tracers of water movement (*Barbeta et al. 2015*). The water use of *Pinus strobus* shifts between deep and surface soil layers under the influence of precipitation (*White et al. 1985*). Research on plants in the Tapares River in the eastern Amazon region of the tropical rainforest found that during the dry season with little precipitation, trees continuously deepened their soil water absorption (*Romero Saltos et al. 2005*). A Study on plants in the riparian forest ecosystem of the lower reaches of the Heihe River found that different water sources contribute differently to plants with groundwater being an important contributor to the growth of desert plants such as *Populus euphratica* (*Zhao et al. 2008*). In the desertification grassland ecosystem of Ordos, simulated rainfall as a single water source showed that the attenuation period of precipitation in typical desert plant water is 7 days (*Cheng et al. 2006*).

Plant water use efficiency is a comprehensive physiological and ecological indicator for evaluating the suitability of plant growth. It reflects the relationship between plant water consumption and dry matter production. It is one of the basic and important characteristics of plants in respond to arid environments, enabling plants to maintain the availability of soil water and thereby improving their resilience to drought conditions (*Hu et al., 2009*). The stable carbon isotope method is based on the stable carbon isotope composition of plants and is currently internationally recognized as an acceptable method for determining long-term plant water use efficiency (WUE). Research has shown a strong correlation between WUE and the stable carbon isotope ratio δ^13^C value. The higher the δ^13^C value of the plant, the higher its WUE, indicating a more economical and conservative water utilization mode (*Shen et al., 2017; Farquhar et al., 1989;Dawson et al., 2002*)

*Cupressus gigantea* is an endemic species along the banks of the Yarlung Zangbo River in the Tibet Autonomous Region, and is also a nationally protected wild plant. Currently, its natural distribution range is known to be limited to both sides of the Yarlung Zangbo River from Jiacha County to Danniang Township, Nyingchi City, and Jubai Conservation Park on the bank of Niyang River which is one of important tributary of Yarlung Zangbo River. Although the existing distribution range is narrow, the climate zone within its distribution range changes from semi-arid to semi humid area. The largest individual of *C. gigantea* has been recorded being more than 2500 years old, with a height of about 55 m, a diameter at breast height of 2.3 m, and a single plant volume of up to 230 m^3^ (*Chen, 1995; Fu, 1995*). *Cupressus gigantea* is an invaluable ecological asset for the environment and climate of the Qinghai and Tibet Plateau, and is also a unique feature of the Yarlung Zangbo River landscape. Its distribution is scattered or clustered in patches. Without proper protection practice, any damage to the population or blockage of regeneration could result in significant losses. Water is an critical factor limiting the growth and distribution of plants in arid and semi-arid areas. Plants can adapt to environmental changes by adjusting their sources of water absorption according to different environmental conditions (*Yang et al., 2015; Gross et al., 2017; José et al., 2007; Barbeta et al., 2015*). In arid environments, plant growth is generally limited by the temporal and spatial availability of water and nutrients (*Veneklaas & Poot, 2003; Alessio et al., 2004* ; *Donovan et al., 2007* ; *Antunes et al., 2018*). The functional traits and population growth of plants in arid and semi-arid ecosystems largely depend on the availability of water in the environment and the plant’s ability to utilize that water (*Palacio et al., 2017*). Therefore, this study explores the following scientific questions: (1) Are there significant differences in soil characteristics across different habitats? (2) Doses *C. gigantea*, which grows in arid environments, exhibit a stronger ability to regulate water use efficiency? (3) Does *C. gigantea* utilizes different sources of water at various stage of growing season?

## MATERIAL AND METHOD

### STUDY AREA

The research site locates in Zhaxitang Village (Fig. 1a) in Lang County, along the Yarlung Zangbo River, Xizang, China. This area has a typical plateau temperate semiarid and semi humid monsoon climate, and the vegetation is *C. gigantea* sparse forest. There is a large temperature difference between day and night. The average annual temperature is above 2°, with the highest temperature in July reaching 17.9 ° and the lowest temperature in January dropping to -12 °. The annual average sunshine hours are 1600-1900 hours. The average annual precipitation is about 500 mm, concentrated from June to September, accounting for 75% to 88% of the total annual rainfall. The maximum rainfall typically occurs from July to August, while the minimum rainfall occurs from November to February. The annual evaporation is 1201 mm (Figure 1b). The soil consists of brown sandy loam with a lot of gravel, characterized by a light texture, low bulk density, weak water storage capacity, and low water content. The climate zone is semi-arid, with a harsh ecological environment and sparse vegetation. *Cupressus gigantea* trees are distributed along the riverside, extending towards the slope direction and up to the upper and middle parts of the mountain. The distribution habitat of *C. gigantea* can be roughly divided into riverbank, terrace, mountain slopes and steep cliff (Fig1.b). This study selected three typical habitats for investigation: the riverside, terrace, and the mountain slope, each with varying vegetation composition (Table 1). A sample plot measuring 100 m×100 m (length ×width) was established within each habitat. The tree height, diameter at breast height, and crown width of each *C. gigantea* individual were measured. Concurrently, soil, plant leaf, and stem samples were collected for analysis of carbon isotopes, physiological, and biochemical indicators. The samples were then transported to the lab and stored in a 4° refrigerator for water extraction to measure hydrogen and oxygen isotope. Sampling occurred at four: August 2022 (rainy season), May 2023 (dry season), August 2023 (rainy season), and November 2023 (before winter) (Fig. 2).

**Table 1.**
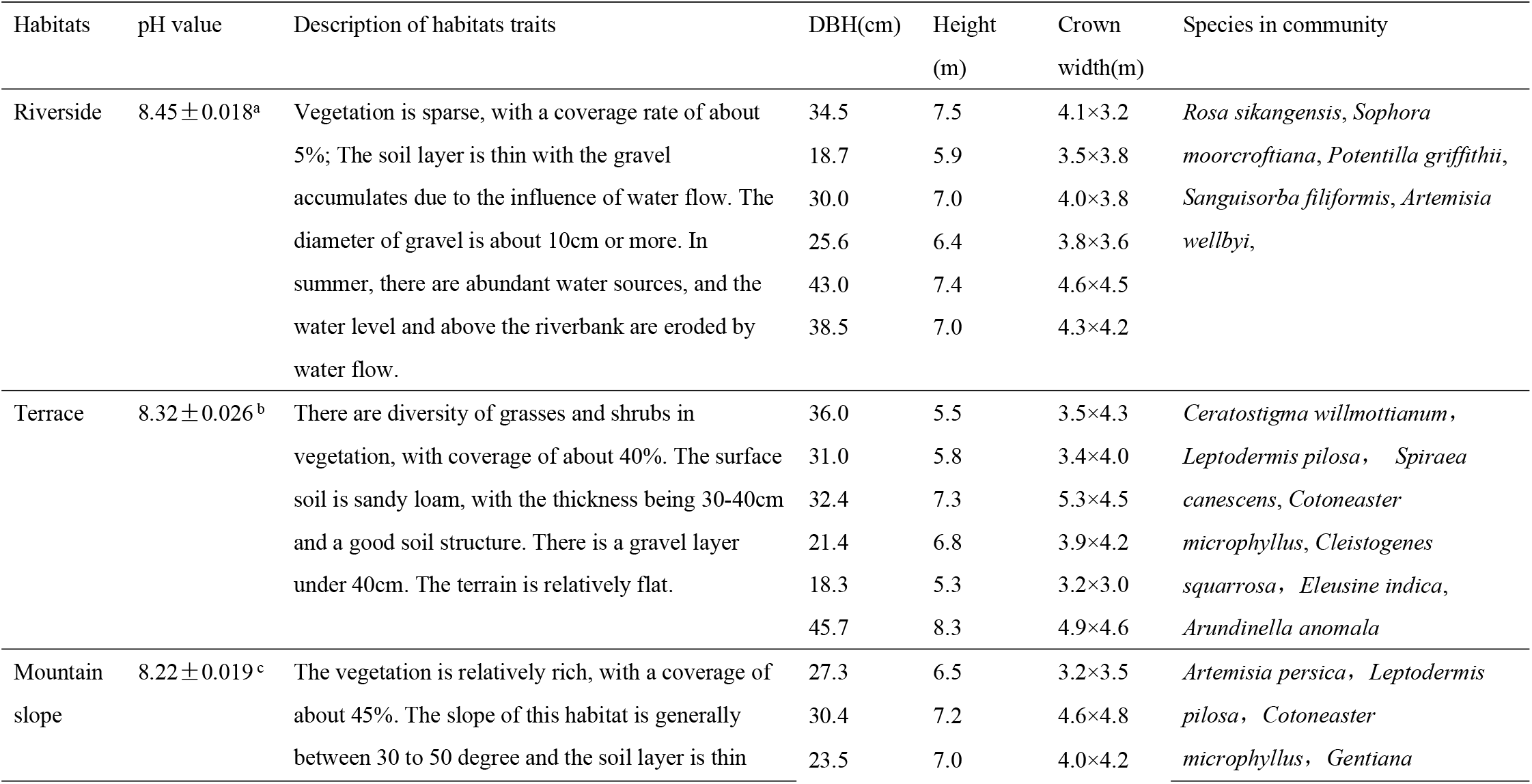
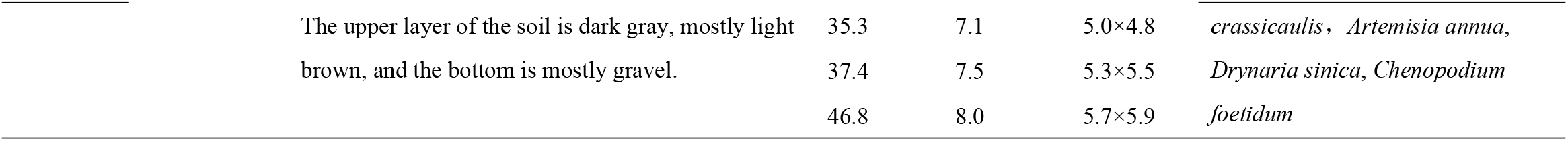
Description of three habitats in pH value of soil layer (0-40cm), habitats characteristics, diameter at breast height(DBH), height of and crown width of *C. gigantea* individuals involved in current experiments and the species compound of plant community at three habitats in Tibet, China. Different lowercase of pH value indicates the significant difference among habitats (*P*<0.05).

**Figure 1.**
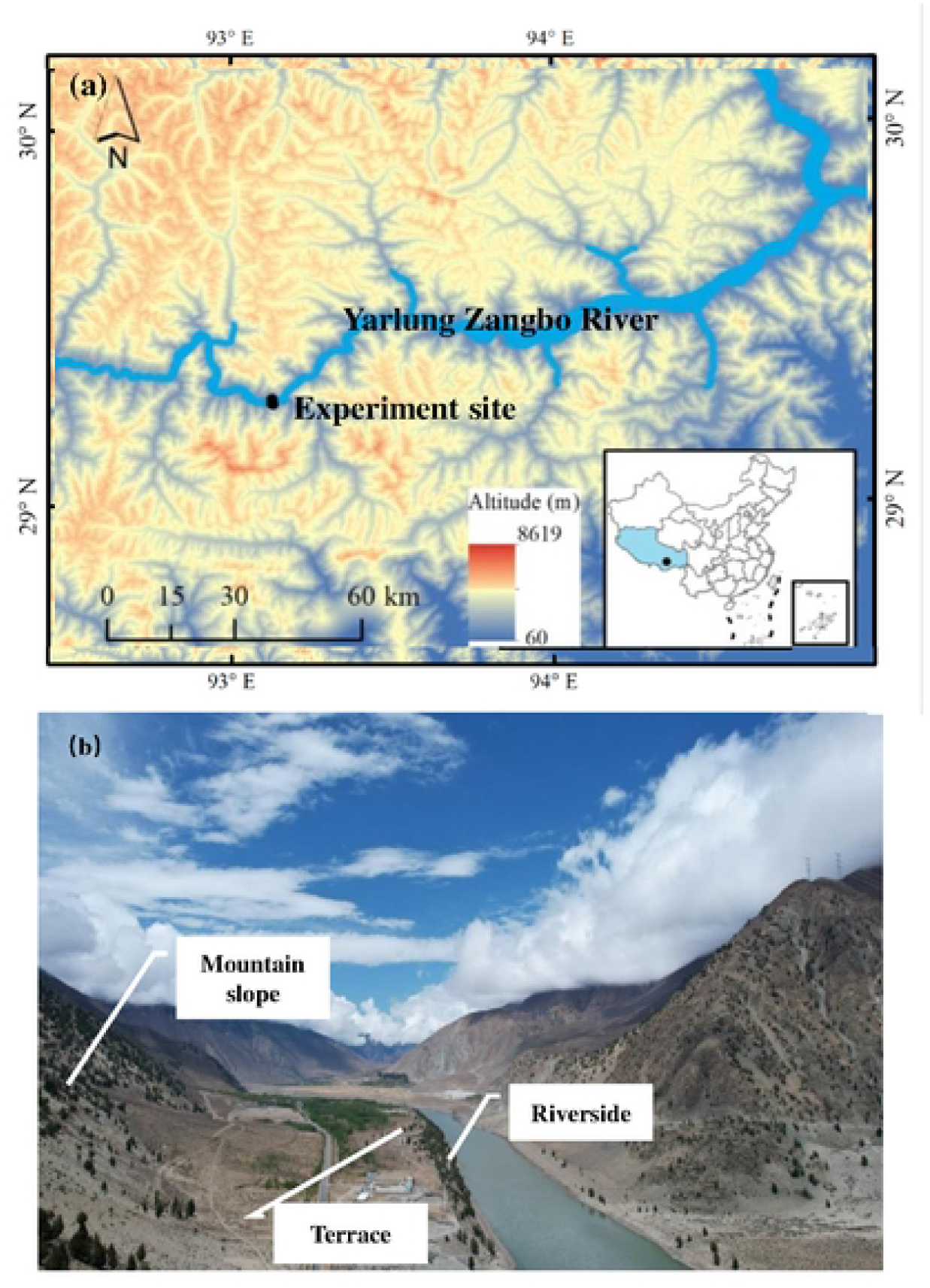
Location of the study site in Zhaxitang village of Lang County, Tibet, China (a) and the distribution of three habitats of *Cupressus gigantea* involved in current experiments (b).

**Figure 2.**
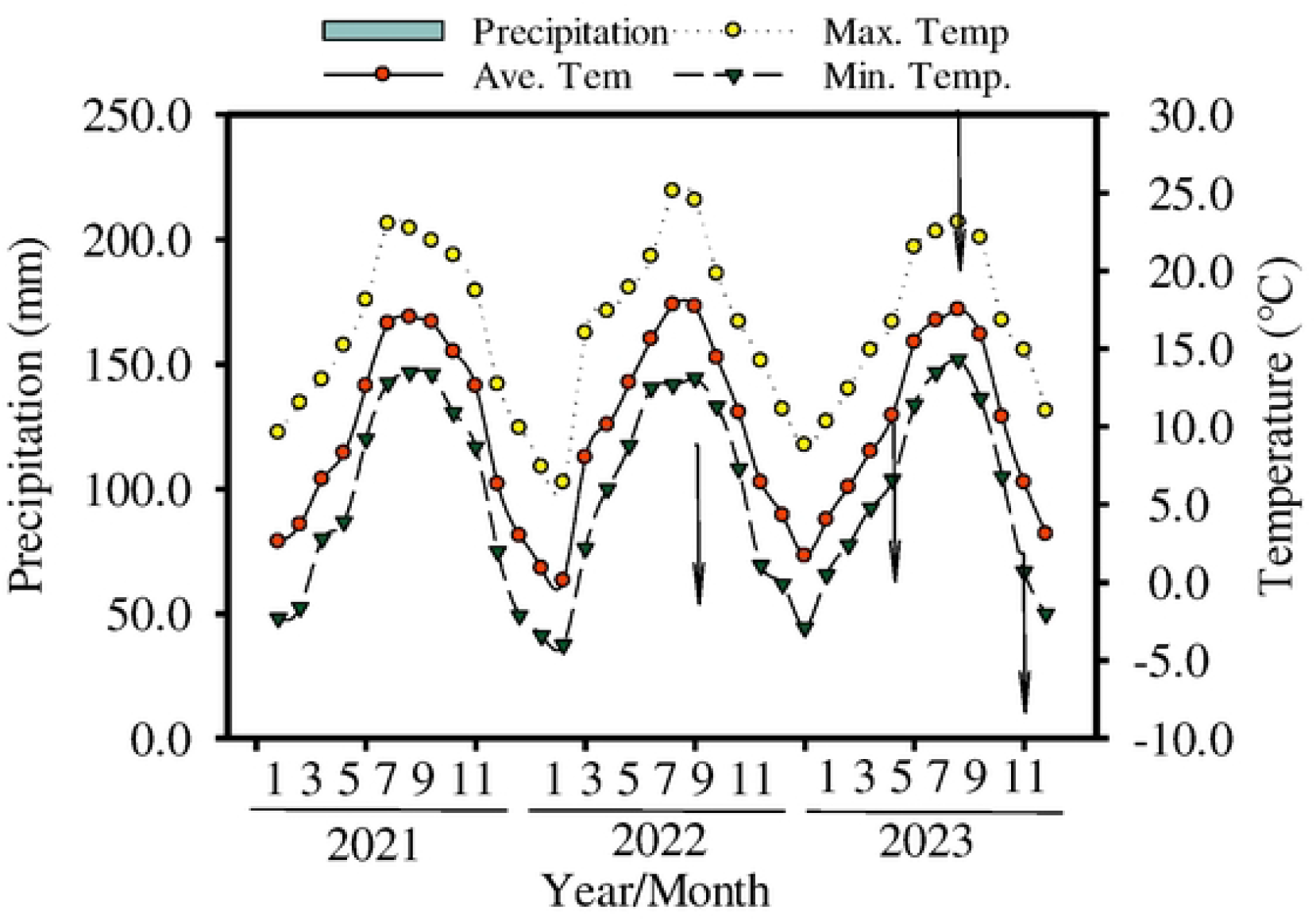
Changes in rainfall (mm), monthly mean atmospheric temperature (°C), monthly maximum and minimum temperatures (°C) during 2021 to 2023. Arrow means the month when the experiments were conducted in Aug.2022, May 2023, Aug. 2023, Nov. 2023 respectively.

### SAMPLE COLLECTION

#### PLANTS AND SOIL SAMPLES

Three groups of *Cupressus gigantea* individual were selected based on a diameter at breast height: 1-20 cm, 20-40 cm, and greater than 40cm. Leaves, branches and roots samples were collected from six individuals in each group. High branches with sufficient light in the middle of the crown were chosen for collecting leaf and branches samples for isotope analysis, water content, and other indicators. To prevent isotope fractionation, the phloem of the branches was removed, and the samples were quickly placed into 8ml glass sample bottles with transparent threaded openings (CNW Technology, Germany). The bottle was labeled and sealed with Parafilm sealing film (American National Can, USA) before being refrigerated and stored at 4° in the laboratory for subsequent water extraction and analysis of hydrogen and oxygen isotopes. Research has reported that the root system of *C. gigantea* is relatively shallow, with the root occupying 73.06% of the vertical range of 0-40 cm (*Bian et al., 2017*). This may be related to the fact that the average soil layer thickness in the natural distribution area of *C. gigantea* is only 30-40 cm. Therefore, this study collected soil samples from 0-20 cm and 20-40 cm soil layers to examine the absorption and utilization of water sources by *C. gigantea*. Soil samples were collected from 50-100 cm away from the trunk in different directions. The samples were quickly placed into transparent threaded 10 ml glass sample bottles, labeled, sealed with a sealing film, and then refrigerated and stored at 4° in the laboratory for subsequent water extraction and analysis of hydrogen and oxygen isotopes. Three soil samples were collected for each selected *C. gigantea* tree. Additionally, soil samples were collected from each habitat to measure physiological and biochemical indicators. Three leaf samples from each diameter class of *C. gigantea* were collected, weighed fresh, and then heated at 105°C for 30 minutes. The samples were subsequently dried at 60 ° C for 48 hours until they reached a constant weight. Weight changes were recorded, and the water content of the leaves was calculated. After drying, the samples were ground, crushed, and sieved through an 80-mesh sieve for carbon isotope determination. For the measuring stable carbon isotope, leaf samples were collected only in August 2022 due to the relatively stable δ^13^C values.

#### WATER SAMPLES

Water samples from the Yarlung Zangbo River were collected during each sampling period near the research site, placed in glass bottles, seal with sealing film, and refrigerate for storage. Three replicates were collected for each sampling period. Underground water samples were collected from the well at Zhaxitang village, with three replicates for each sampling period. Rainwater samples were collected during the sampling period, with at least three rainfall events collected from August to September 2022, May to June 2023, July to August 2023, and October to November 2023. All samples are sealed in 8ml glass sample bottles, labeled with the year, month, and day, refrigerated, and transported back to the laboratory for storage at 4 ° until measurement.

### SAMPLES ANALYSIS

#### MEASUREMENT ISOTOPE AND WATER SOURCE DETERMINTAITON

Water was extracted from *C. gigantea* branches and soil samples using a low-temperature vacuum extraction method to prevent isotope fractionation during the extraction process. The hydrogen and oxygen stable isotope analysis of both extracted water samples and field-collected water samples was performed using a stable isotope ratio mass spectrometer (253 plus) to determine the δD and δ^18^O values. The experimental results were corrected using the Vienna Standard Mean Ocean Water (VSMOW), and the measured isotope results were reported as the the difference (‰) relative to VSMOW standard value. The stable isotope ratio is expressed as follows:

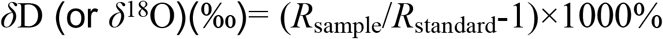

*R*_sample_:isotope ratio of samples;*R*_standard_:Isotope ratio of standard materials.

The multi-source linear mixed model (IsoSource) determines the total contribution rate of all sources by summing to 100%, and then calculates the upper and lower limits of the contribution rate error for each source. The procedure involves inputting potential source waters into the IsoSource software: soil water (0-20cm), soil water (20-40cm), groundwater, and river water. Next, isotopic values and markers from the *C. gigantea* plant samples are entered. After setting the increment and allowable deviation, the program is run to perform the calculations and save the results.

IsoSource The multiple linear model is as follows:

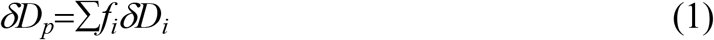

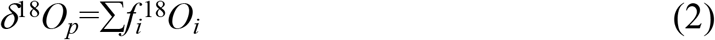

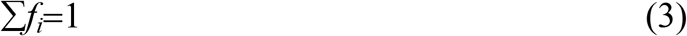

*δ*D_i_(*δ*^18^O_i_):Hydrogen and oxygen isotope values of each water source; *δ*D_p_(*δ*^18^O_p_):Abundance of hydrogen and oxygen isotopes in plant water;*f*_i_: Contribution rate of water source i to plant water content.

This study includes four potential water sources: soil water (0-20cm), soil water (20-40cm), groundwater, and river water. Therefore, the multiple linear model used in this study is:

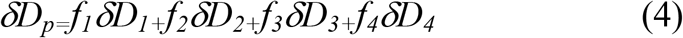

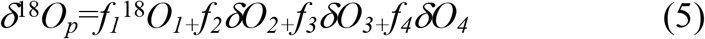

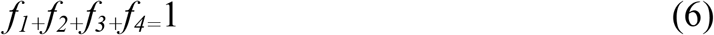

The contribution rates of four water sources to the water content in the xylem of *C. gigantea* are represented as following: *f*_1_, soil water (0-20 cm), *f*_2_, soil water (20-40 cm), *f*_*3*_, groundwater, and *f*_4_, river water.

#### CARBON ISOTOPE DETERMINATION WATER USE EFFICIENCY

Carbon isotopes of leaf samples were analyzed using a stable isotope mass spectrometer (Delta plus xp). Calibrated CO_2_ was used as the standard for determining the stable carbon isotope ratio, with international standard V-PDB serving as the reference gas.

The sample was compared to the standard gas to calculate the *δ*^13^C value of the *C. gigantea* leaves. This value represents the isotope ratio relative to the international standard. The average water use efficiency (WUE) and leaf isotope resolution *Δ* can be calculated using a linear model using the following formula:

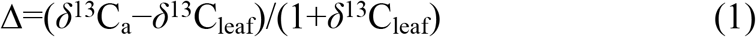

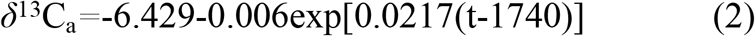

*δ*^13^C_leaf_ is the carbon isotope ratio of plant leaves, *δ*^13^C_a_ is the carbon isotope ratio of environmental CO_2_, which is -6.7‰ (Yue et al., 2022); t is the sampling year.

Therefore, the leaf isotope resolution (*Δ*) can be calculated using the formula above. For this study, with the sampling year being 2022, the calculation is as follows:

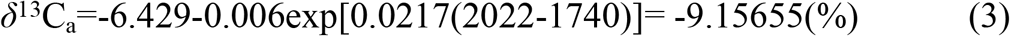

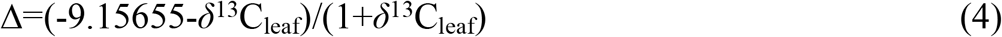

The *C*_i_/*C*_a_ ratio is an important physiological and ecological indicator of plants, reflecting the relative amount of net assimilation rate and stomatal conductance in relation to CO_2_ demand and supply. The calculation formula is as follows (Huang et al., 2019):

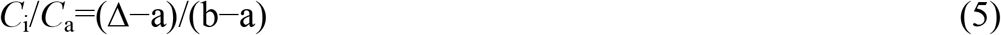

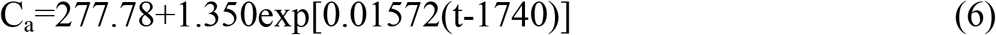

*C*_i_ is the intercellular CO_2_ concentration; *C*_a_ is the concentration of environmental CO_2_ in the study area (μmol mol^-1^); *a* is the isotopic fractionation coefficient (4.4%) caused by the diffusion of environmental CO_2_ in still air; b is the fractionation coefficient (27%) resulting from CO_2_ fixation in chloroplasts and its internal diffusion through 1,5-bisphosphoribulose carboxylase (Rubisco); t is the sampling year.

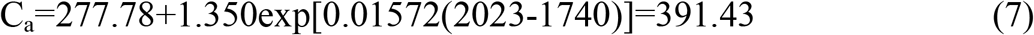

The formula for calculating the average water use efficiency (WUE) is as follows:

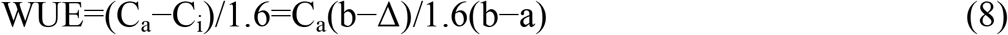

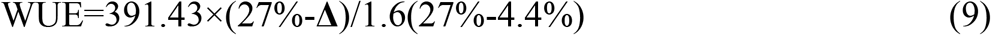

#### DETERMINATION OF BIOCHEMICAL INDICATORS FOR SOIL AND LEAVE

Leaf and soil indicators were measured at different habitat sampling sites in August 2022. The physical and chemical properties of soil and *C. gigantea* leaves were determined using 10 indicators: soil pH value, soil total nitrogen, soil ammonium nitrogen, soil nitrate nitrogen, soil available phosphorus, soil organic carbon, leaf moisture content, leaf total nitrogen, and leaf total carbon. The soil pH value was measured using an acidity meter; Leaf moisture content was determined by the drying method; Total nitrogen content in soil and leaves was measured using the Kjeldahl nitrogen determination method; Organic carbon content in soil and total carbon content in leaves were measured using the potassium dichromate external heating method; Soil ammonium nitrogen and soil nitrate nitrogen content were determined using the flow analysis method; Soil available phosphorus content was measured using the NaHCO_3_ extraction molybdenum antimony colorimetric method.

#### DATA ANALYSIS

The data were organized and preliminarily processed using Excel 2019, and analyzed using SPSS 25.0 and R 4.3.1 software. One-way analysis of variance (ANOVA) and least significant difference (LSD) method were employed. The LSD analysis compared differences in stable oxygen isotope ratios and physicochemical indicators of soil water sources at various sampling periods, with a significance level set at p=0.05. Graphic were created using R4.3.1, Origin 2021,, ArcGIS 10.8, and Adobe Illustrator 2022.

## RESULTS

### TRAIT OF PLANT AND SOIL AMONG HABITATS

The altitude of the three habitats increases progressively from the riverside to the terrace and then to the mountain slope (Table 1). Soil total nitrogen (TN_soil_), soil organic carbon content (SOC), soil nitrate nitrogen (NO_3_-N), and ammonium nitrogen (NH_4_-N) were significantly higher at the mountain slope compared to the other two habitats (Fig. 3a, b,c,d) (*P*<0.05). The soil pH value was significantly higher at the riverside habitats than at the other two habitats (*P*<0.05), with no significant difference in pH values between the terrace and mountain slope habitats (Table 1). The total carbon content in the leaves was significantly higher at the terrace compared to the riverbank and mountain slope (*P*<0.05) (Fig. 4a) ; The total nitrogen in the leaves was the highest at riverside habitats, the lowest at mountain slope habitats, and differed significantly among three habitats (*P*<0.05) (Fig. 4b). The water content of *C. gigantea* leaves was the lowest at mountain slope habitats compared to the other two habitats (*P*<0.05), with no significant difference between the riverbanks and terrace habitats (Fig. 4c).

**Figure 3.**
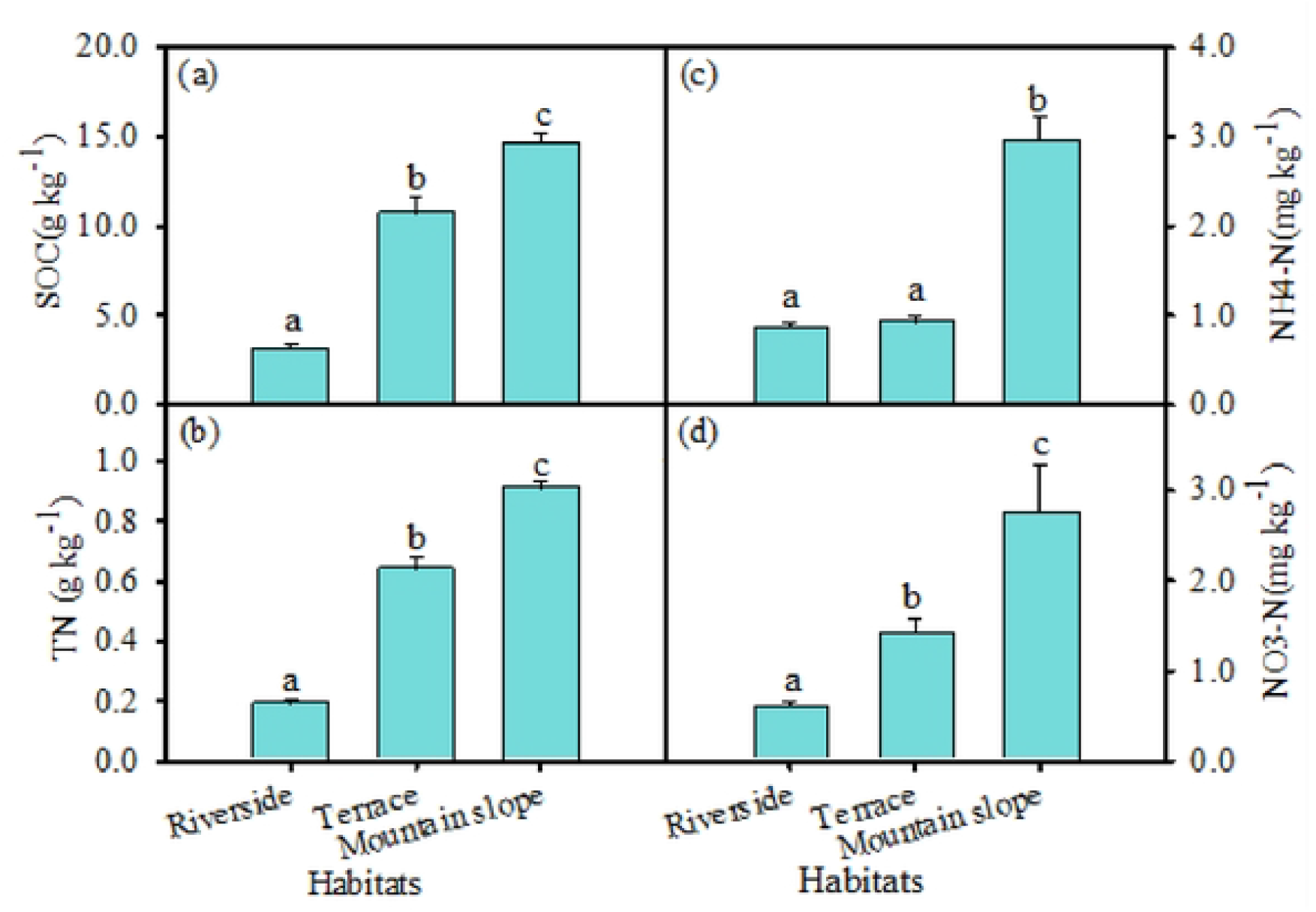
Summary of soil organic carbon (SOC) (a), soil total nitrogen (TN) (b), NH4-N (c) and N03-N (d) of soil (0-40cm) from Riverside, Terrace and Mountain slope carbon in Aug. 2022.Values are means of six replicates (+SE). Different lowercase letters indicate significant difference among habitats (P<0.05), same letters mean no significant different (P>0.05).

**Figure 4.**
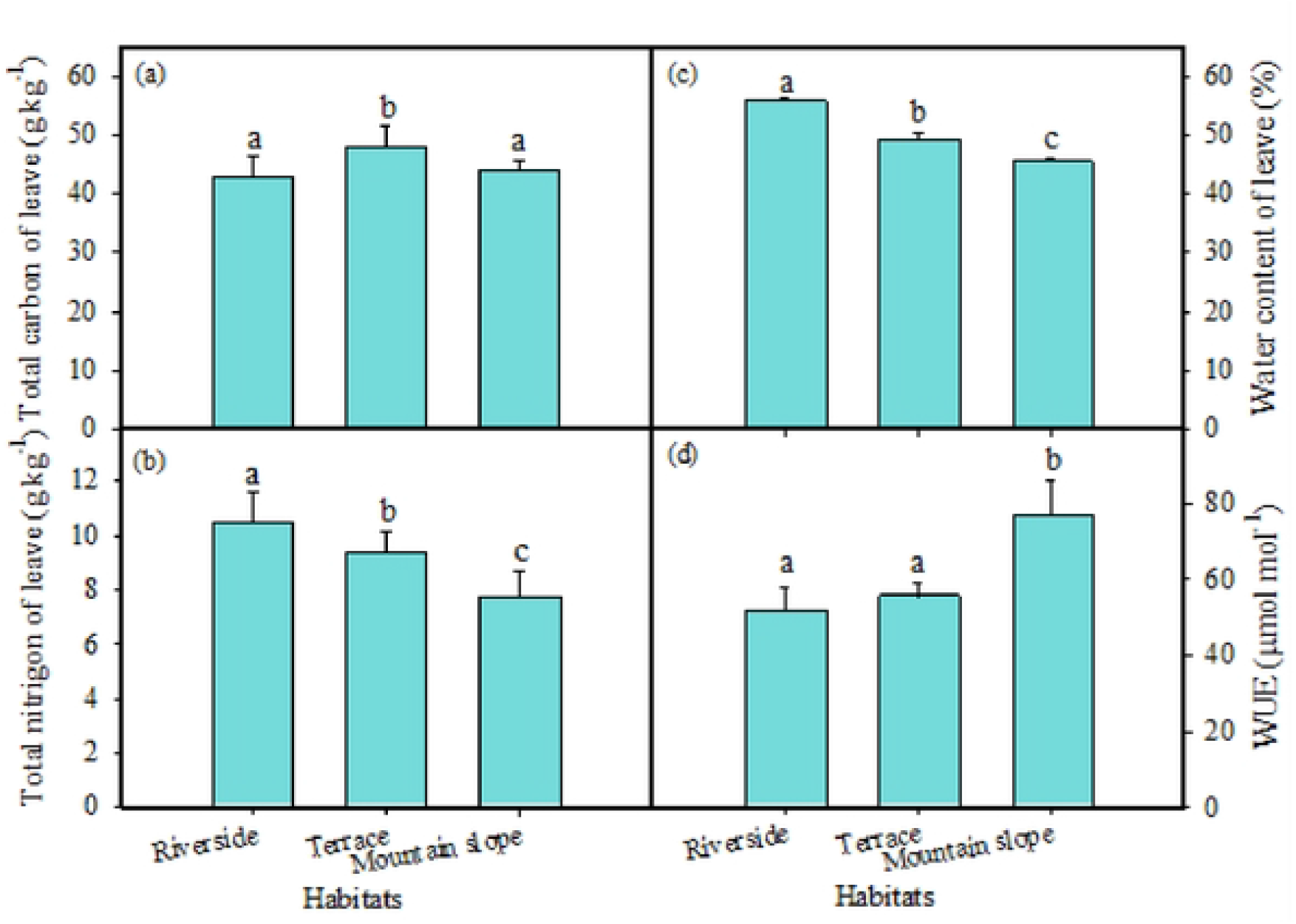
Summary of total carbon (TC_leaf_) (a), total nitrogen (TN_leaf_) (b), water content of leaf(c) and water use efficiency (WUE) (d) of plant leave Crom Riverside, Terrace and Mountain slope in Aug. 2022. Values are means of six replicates (+SE). Different lowercase letters indicate significant difference among habitats (P<0.05), same letters mean no significant different (P>0.05).

The δ^13^C values in *C. gigantea* leaves were significantly higher at the mountain slope compared to the other two habitats (*P*<0.05) (Table 2). The average δ^13^C value at the mountain slope was -23.4 ‰±0.67, with a range of -24.1 ‰ to -22.4 ‰. The δ^13^C values of the riverside and terrace habitats were -25.1 ‰±0.98 and -25.0 ‰±1.01 respectively. Water use efficiency (WUE) was calculated based on the δ^13^C value of plant leaves (*Farquhar et al., 1982; Marshall & Zhang, 1994*). WUE was significantly higher at the mountain slope, with the highest average values of 98.3 μmol mol^-1^(*P*<0.05). The WUE of *C. gigantea* was similar at the riverbank and terrace habitats, with an average value of 63.5 μmol mol^-1^ and 64.2 μmol mol^-1^, respectively (Fig.4d). WUE showed a highly positive correlation with soil organic carbon (SOC),soil total nitrogen (TN_soil_), soil NH4-N, and soil pH values (*P*<0.01), and a highly significant negative correlation (*P*<0.01) with leaf water content (LWC) (*P*<0.01) (Fig.5).

**Figure 5.**
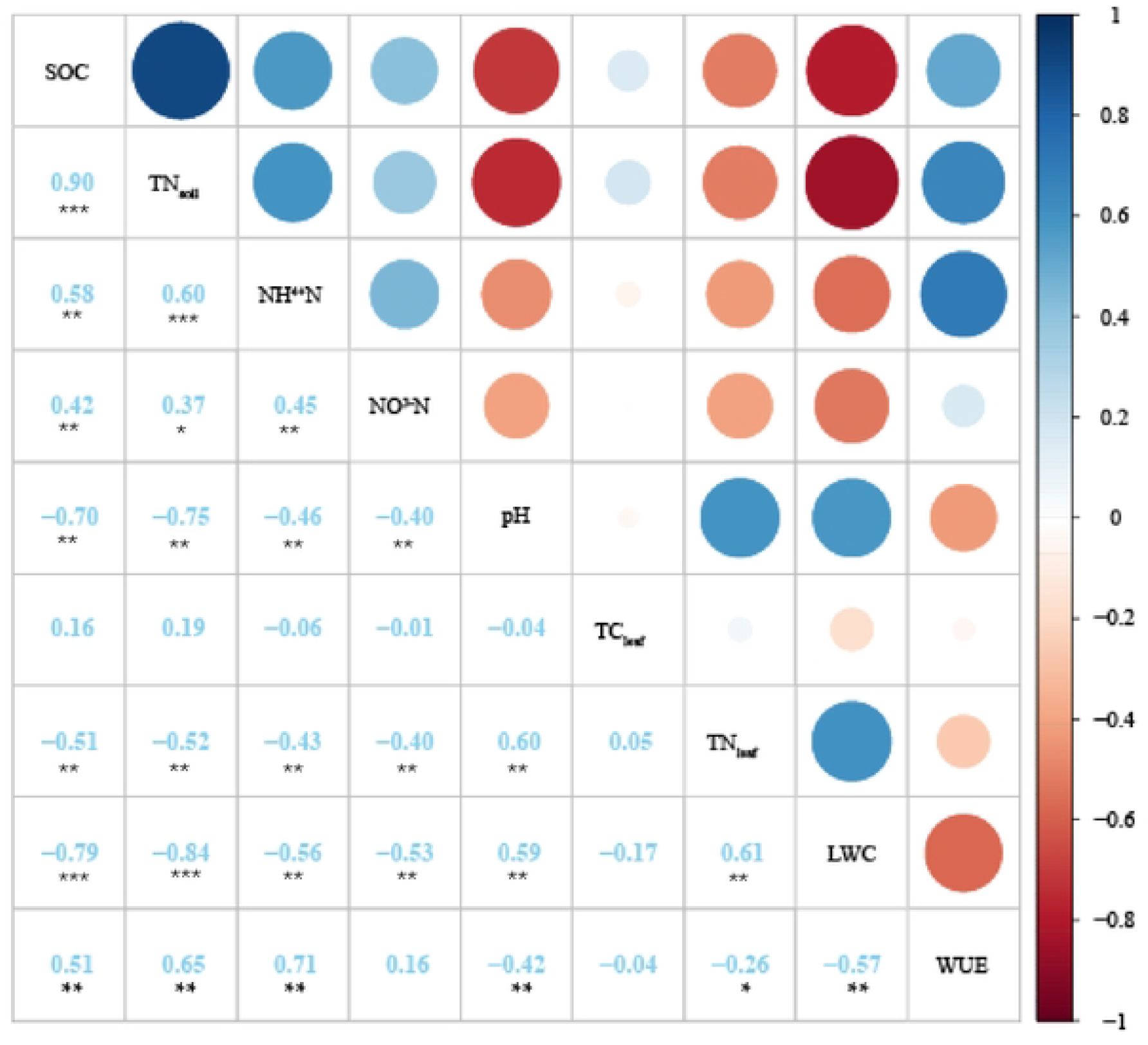
The relationship of water use efficiency of *Cupressus giganrea* in relation to soil organic carbon (SOC), soil total nitrogen (TN_soil_), NH4-N, N03-N of soil (0-40cm), total carbon (TC_leaf_), total nitrogen (TN_leaf_), water content of leaf. * P<0.05; ** P<0.01; ***P<0.00 1.

**Table 2.** Results of linear mixed-effects model analysis (F-values, degree of freedom and significance) testing the effects of growing seasons, habitats, soil layers and their interactions on *δ*D, *δ*^18^O values of soil and plant leaves of *C. gigantea*. Data were performed on log_e_-transformed values. Significant level; \**P*<0.05; \*\**P*<0.01; \*\*\**P*<0.001.

| Variables | $\delta D_{\text{soil}}$ | $\delta^{18}O_{\text{soil}}$ | $\delta D_{\text{plant}}$ | $\delta^{18}O_{\text{plant}}$ |
| --- | --- | --- | --- | --- |
| Seasons | 71.23(3,120)*** | 21.31(3, 120)*** | 53.02(3,60)*** | 23.95(3,60)*** |
| Habitats | 5.41(2, 120)** | 7.52(2, 120)*** | 41.78(2,60)*** | 18.96(2,60)*** |
| Soil layers | 41.53(1, 120)*** | 0.38(1, 120) | ----- | ----- |
| Season × habitats | 32.25(6, 120)*** | 7.82(6, 120)*** | 11.85(6,60)*** | 4.21(6,60)*** |
| Seasons × soil layers | 13.59(2, 120)*** | 2.39(3, 120) | ----- | ----- |
| Habitats × soil layers | 13.85(3, 120)*** | 3.78(2, 120)* | ----- | ----- |
| Seasons × habitats × soil layers | 15.30(6, 120)*** | 6.19(6, 120)*** | ----- | ----- |

**Table 3.** Results of two-way ANOVA analysis (F-values, degree of freedom and significance) of the effects of seasons, habitats and their interactions on leaf water content (LWC), contribution of soil water (0-20cm) to plant, and (20-40cm) *δ*^18^O values of soil and plant. Data were performed on log_e_-transformed values. Significant level: \**P*<0.05; \*\**P*<0.01; \*\*\**P*<0.001.

| Variables | Leave water content | Water contribution of soil layer (0-20cm) | Water contribution of soil layer (20-40cm) | Water contribution of ground water | Water contribution of river water |
| --- | --- | --- | --- | --- | --- |
| Seasons | 21.62(3,60)*** | 6.15(3,60)*** | 6.76(3,60)*** | 1.58(3,60) ns | 2.41(3,60) ns |
| Habitats | 44.38(2,60)*** | 71.40(2,60)*** | 24.29(2,60)*** | 3.694(2,60)* | 28.74(2,60)*** |
| Seasons × habitats | 2.26(6,60)* | 12.16(6,60)*** | 6.15(6,60)*** | 3.62(6,60)** | 3.36(6,60)** |

### THE *δ*D AND *δ*^18^O VALUES FROM DIFFERENT WATER SOURCES

The *δ*D and *δ*^18^O values of rainfall changed significantly over the experiment period, being most enriched in May 2023, with average value of -46.1 ‰ and -7.2 ‰, respectively. These values were the most impoverished in August, 2022, with an average value of -103.3 ‰ and -13.6 ‰, respectively (Fig. 6). The slope of the Local Meteorological Water Line (LMWL) is slightly lower than that of the Global Atmospheric Waterfall Line (GMWL: *δ*D=8 *δ*^18^O+10), which is a more significant trends in arid regions (Fig. 6). This is mainly due to the influence of secondary evaporation on the precipitation process, where lighter *δ*D is depleted, and heavier *δ*^18^O is enriched in precipitation. During the study period, the average values of *δ*D and *δ*^18^O in groundwater were -110.1 ‰ and -14.6 ‰ respectively, which were distributed slightly lower and to the right of the atmospheric precipitation line. The average values of *δ*D and *δ*^18^O of the Yarlung Zangbo River water were -121.0 ‰ and -16.4 ‰ respectively, which were distributed on both sides of the local atmospheric precipitation line (Fig. 6). The average values of soil water *δ*D and *δ*^18^O are -92.5‰ and -7.0 ‰, respectively (Fig. 6), distributed in the lower right corner of the local atmospheric precipitation line, indicating that precipitation undergoes significant evaporation, enrichment, and fractionation after entering the soil. Due to the influence of evaporation degree and supply source, there are significant differences in the slope and intercept of each water body, but all are smaller than the slope of the regional atmospheric precipitation line, showing varying degrees of deviation. This indicates that each water body is supplied by precipitation.

**Figure 6.**
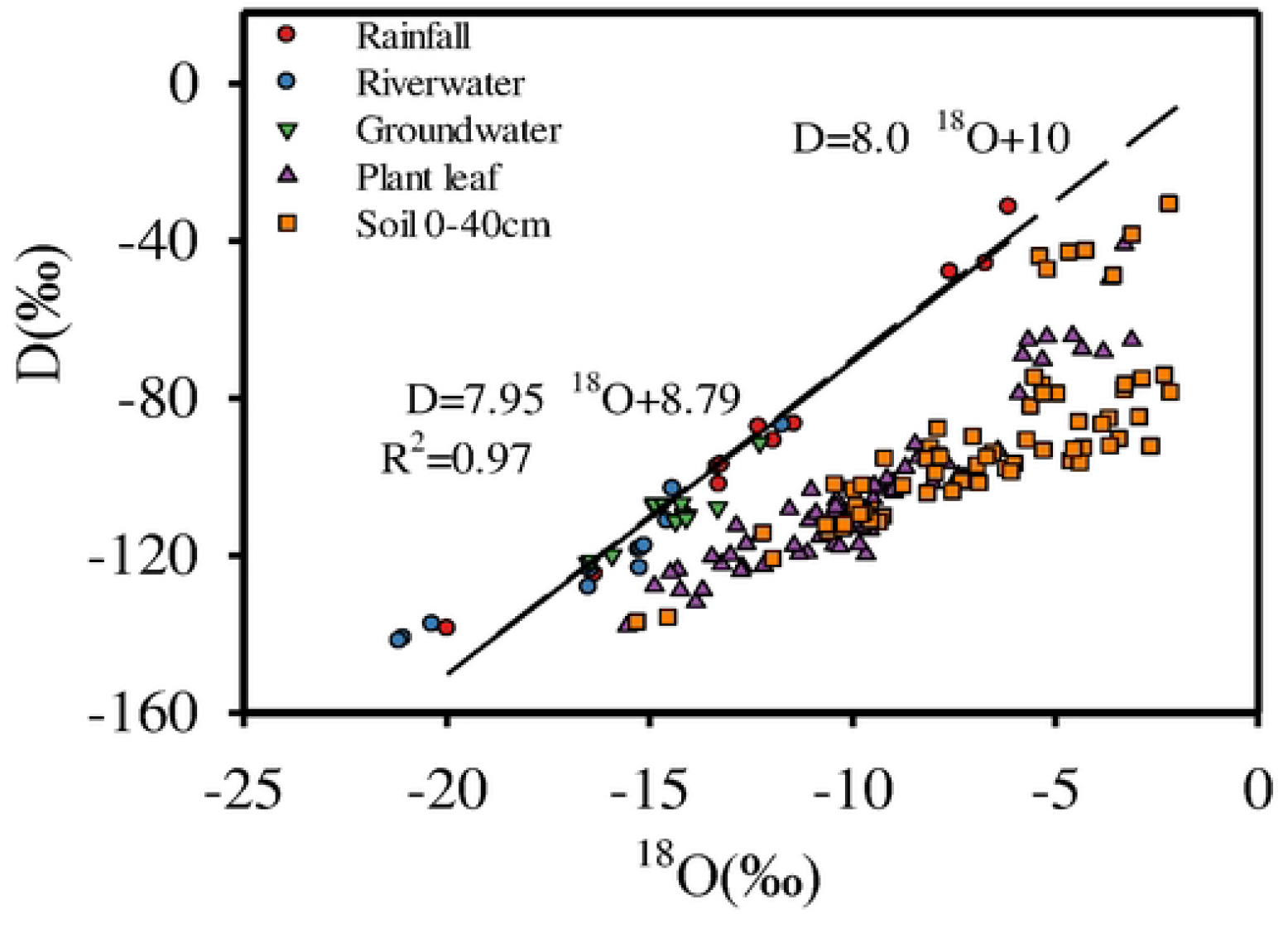
Stable isotopic compositions of rainfall, river water, groundwater, soil water and plant water at the study site. The local meteoric water line is very close to the GMWLwhose equation is δD = 8.0 ×δ^18^O+ 10.

### VARIATION OF *δ*D AND *δ*^18^O VALUES IN DIFFERENT WATER SOURCES

The average *δ*D and *δ*^18^O values of rainfall in May 2023 were the highest, being -46.1 ‰ and -7.2 ‰, respectively, while they were the lowest in Nov. 2023, being -125.9 ‰ and 16.7 ‰ respectively. The change of rainfall among different seasons was quite significantly (*P*<0.001). The average values of *δ*D and *δ*^18^O in river water in Aug. 2023 were significantly lower than other seasons, being -140.1 ‰ and -20.9 ‰, respectively. However, they were similar between Aug.2022 and May 2023, Nov. 2023 due to the less rainfall in Aug. 2022. This trend indicates that during the sampling period of the study, the changes in the values of *δ*D and *δ*^18^O in the river water displayed a significant trend of impoverishment during the rainy season and gradual enrichment during the non-rainy season. The range of *δ*D and *δ*^18^O value in groundwater are not large, and relatively stable, with average values being -110.1 ‰ and 14.6 ‰, respectively. The average values of *δ*D and *δ*^18^O in groundwater were greater than those of river water, indicating that river water is the main source of groundwater.

The average *δ*D and *δ*^18^O values of soil water at different periods in the study area showed significant differences (*P*<0.05) (Table 2 and Fig.7b,c,e,f). The *δ*D and *δ*^18^O value of surface soil layer (0-20cm) at Mountain slope habitats were higher than those at the other two habitats, but no significance differences were observed in the lower soil layer (20-40cm) (Fig.7b,c,e,f). From a seasonal perspective, the average values of soil water were the lowest in Aug. 2023 and the highest in May 2023.

**Figure 7.**
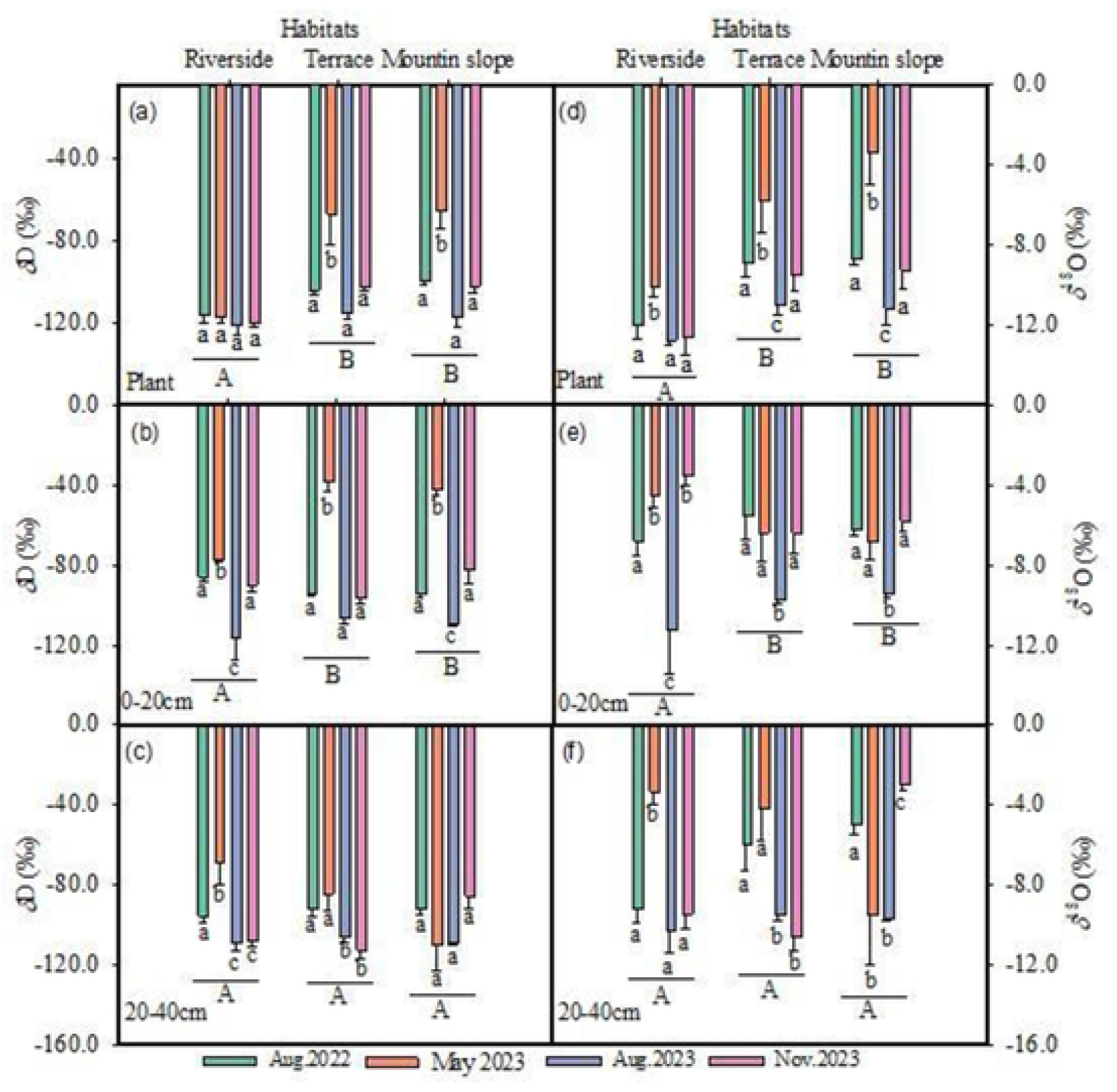
The changes of δD values and δ^18^O values of plant (a,d), soil of 0-20cm layer (b,e), soil of 20-40cm (c,t) at Riverside, Terrace and Mountain slope habitats in four different sampling period. Values are means of four replicates (±SE). Different lowercase letters indicate significant difference among habitats (*P*<0.05), same lowercase letters mean no significant different (*P*>0.05). Different capital letters mean significant difference among habitats (*P*<0.05), same capital letters indicate no difference.

### VALUES OF *δ*D AND *δ*^18^O in XYLEM WATER

The *δ*D values of the xylem water of *C. gigantea* at riverside habitats were significantly lower than those of the terrace and mountain slope habitats (*P*<0.05). However, the values did not change significantly across different seasons, indicating that the water sources of the plants did not switch among seasons in riverside habitats. The average *δ*D and *δ*^18^O values of xylem water were significantly higher in May 2023 than in other seasons, being -67.3 ‰ ± 8.43 and -5.9 ‰ ± 1.49 at Terrace habitats,and 65.62 ‰ ±8.50 and 3.46 ‰ ±1.53 at Mountain slope habitats (Fig.7a, d). The *δ*D and *δ*^18^O values of the xylem water of the *C. gigantea* at mountain slopes showed significant seasonal variations, with highest values in May 2023 and the lowest in Aug. 2023 (*P*<0.01) (Fig.7 a,d). At terrace habitats, the highest value was observed in the early growth season of May 2023, with average *δ*D and *δ*^18^O values of -67.3 ‰ ± 8.43 and -5.9 ‰ ± 1.49, respectively. In August 2023, the values were the lowest, being of -117.7 ‰ ± 4.52 and -11.19 ‰ ± 0.81 (Fig.8 a, d).

### QUANTITATIVE ANALYSIS OF ABSORPTION WATER SOURCES

Using the IsoSource multiple linear mixed model, the contribution rates of potential water sources to *C. gigantea* at different stages in three different habitats of riverside, terrace, and mountain slope were quantified (Fig.6). The results showed significant difference in water sources of *C. gigantea* with the alternation of dry and rainy seasons, and the contribution rates of each potential water source varied significantly among habitats. The utilization ratio of Yarlung Zangbo River water for *C. gigantea* along the riverside was significantly higher than in other habitats except during the rainy season Aug. 2023 (Fig.8a,b,c,d). The utilization ratio of river water for *C. gigantea* at the riverside reached 66.0%, 78.6% and 55.5% in August 2022, May 2023 and November 2023 respectively. In August 2023, the rainfall in the region increased, and the utilization rate of four water sources by *C. gigantea* are similar: 27.7 % for upper soil water (0-20cm), 24.1% for lower soil water (20-40cm), 24.7% for groundwater and 23.6% for river water, soil water became the main source of water for *C. gigantea*, with a total contribution of 51.8% (Fig.8). The rainy season in August 2022 was an exception due to extreme drought conditions with low rainfall, and the main source of absorption for the *C. gigantea* was still river water.

The water use of *C. gigantea* in the mountain slope and terrace habitats was relatively stable across different seasons, with soil water as the main source. The utilization rate of river water was significantly lower than that for *C. gigantea* in the riverside areas. The utilization rate of soil water by *C. gigantea* was similar in May, August and November 2023, with utilization ratios exceeding 50%, making it the main source of water in these regions. The lowest utilization ratio of soil water by *C. gigantea* on the mountain slope occurred in November 2023, while the utilization ratio of river water and groundwater increased. In other periods, the utilization ratio of soil water by *C. gigantea* exceeded 60% (Fig. 8). The utilization rate of groundwater by *C. gigantea* on the mountain slope was lower in August 2023 than in May, August, and November 2023 (Fig. 8c), likely due to the increased rainfall in August 2023.

**Figure 8.**
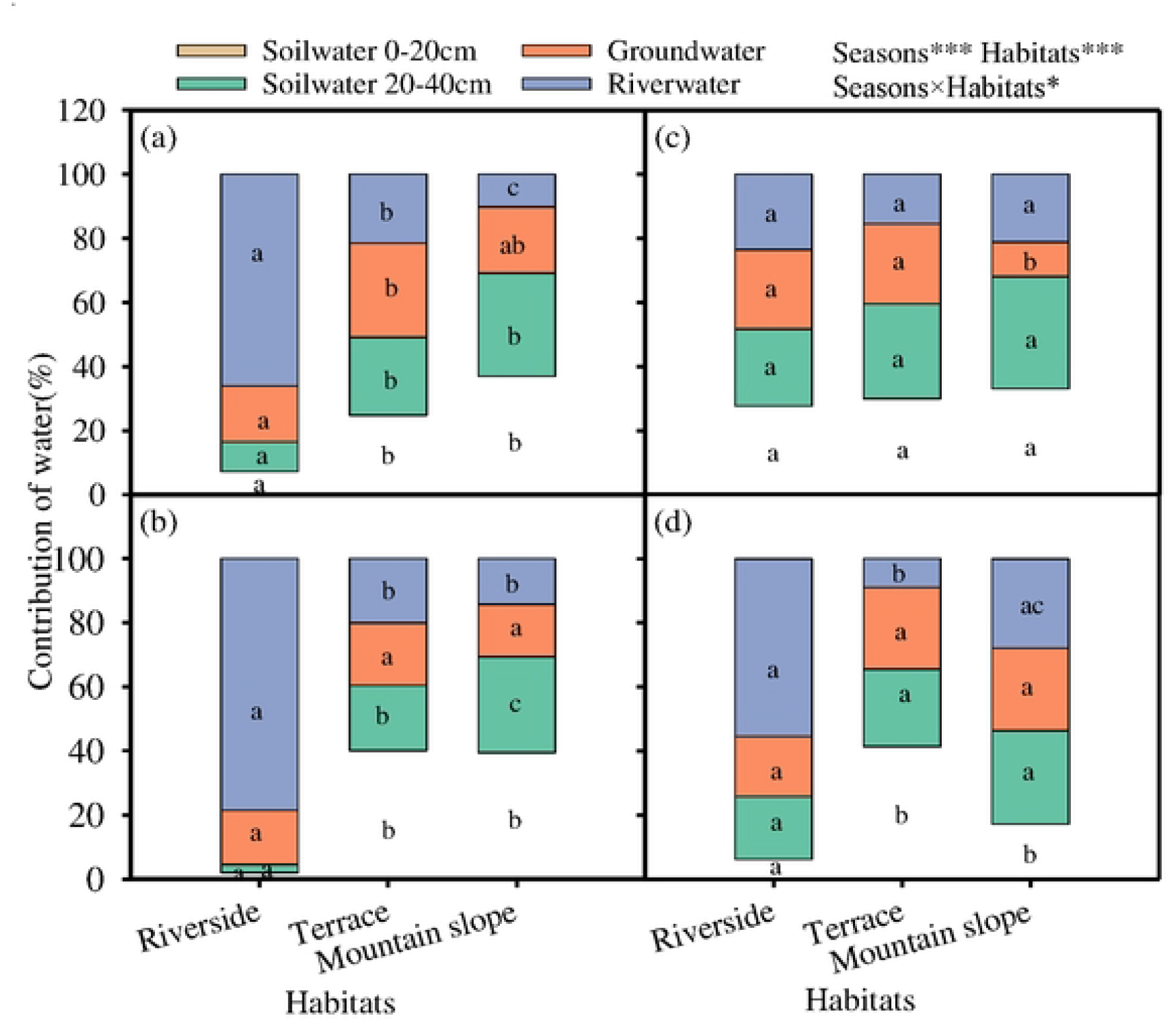
Contribution of four water source, *i*.*e*. soil water (0-20cm), soil water (20-40crn), river water and ground water to the plant at Riverside, Terrace and Mountain slope habitats of study site in Aug. 2022 (a), May 2023 (b), Aug. 2023 c) and Nov. 2023 (d). Different lowercase letters indicate significant difference among habitats (*P*<0.05), same letters mean no difference (*P*>0.05).

## DISCUSSION

### WATER USE EFFICIENCY OF *C. gigantea*

Soil, as a carrier for plant survival, exhibits heterogeneous spatial and temporal distribution of its physicochemical properties (*Campbell & Grime, 1989*). As the main distribution center of *C. gigantea*, Lang County has a harsh ecological environment, steep mountains, sandy soil, sparse vegetation. The habitat heterogeneity and fragmentation are severe in the distribution area of *C. gigantea* (*Luo et al., 2010*). One of the fundamental and important characteristics of plants in responding to stressed environments is their capability to regulate the water use efficiency (*Onoda 2017; Wang et al., 2010*). This enables plants to adapt to changes in soil water availability, thereby enhancing their ability to cope with arid environments (*Su et al. 2020; Hu et al., 2009; Cao et al., 2012*). There are significant differences in the physicochemical properties of soil and *C. gigantea* leaves in different habitats. At mountain slope habitats with high altitude, soil organic carbon, soil nitrate nitrogen, soil ammonium nitrogen, and soil organic carbon were higher than that of terrace and riverside habitats.

This may be due to the decrease in atmospheric and soil temperatures with increasing altitude, which makes it difficult for organic matter to decompose, resulting in an increase in surface total nitrogen content (*Wei et al., 2014*); The soil pH value shows a decreasing trend with increasing altitude, and the soil at terrace and mountain slope habitats was closer to neutral than that at the riverbank.

The total carbon content, leaf water content, and total nitrogen content of *C. gigantea* leaves decrease at mountain slope habitats with increasing altitude, and there are significant differences among different habitats. The leaf water content of plants can significantly affect the intensity of photosynthesis and reflect the water and heat conditions of the surrounding habitat. The decrease in water content of *C. gigantea* leaves with increasing altitude suggests that soil moisture conditions in high-altitude areas are slightly lower than those at the riverbank and Terrace. The decrease in total nitrogen content in leaves may be due to the increased total solar radiation and ultraviolet radiation at high altitudes. Plants need to consume their own energy to accumulate resistance to ultraviolet radiation and avoid damage. Additionally, insufficient available soil water affects plant photosynthesis (*Zhang et al., 2010; Querejata et al. 2021, 2022; Reich et al, 2009*). The changes in leaf nitrogen content and water content are complementary. Habitat differences such as altitude often cause changes in environmental factors such as light, temperature, and rainfall, which in turn affect plant δ^13^C values and water use efficiency (WUE) (*Allen et al., 2010; Wright et al. 2001; Lu et al. 2024*). The leaves of *C. gigantea* in the mountain slope habitats have the highest δ^13^C value (−22.38%), which is significantly different from those in the other two habitats. The leaves of *C. gigantea* in the riverside habitat have the lowest δ^13^C value (−26.56%). The variation in WUE value of *C. gigantea* among different habitats is consistent with the δ^13^C value. The leaves of *C. gigantea* in the mountain slope habitat have the highest WUE being 107.8 μmol mol^-1^, while the leaves in riverside habitats have the lowest WUE being 35.7 μmol mol^-1^. This indicates that the water use efficiency of *C. gigantea* leaves has strong plasticity to cope with changes in habitat conditions. The difference between the δ^13^C and WUE values of plant leaves is influenced by various factors such as temperature, light intensity, and water content in the environment. Plants are capable of perceive external environmental changes and regulate their physiological morphology, such as reducing stomatal conductance, which affects the δ^13^C value of plant leaves, thereby regulating WUE to adapt to environmental changes (*Querejeta et al. 2022; Wen et al., 2010; Song et al., 2011; Wright et al., 2003*). Relevant research shows that the leaf δ^13^C value of *Q. aquifolioides* rises with the elevation above 2800 m in the eastern slope of the Himalayas, Tibet, which is similar to the plateau environment of the current study (Li et al., 2006). Additionally, studies on two species, *Hippophae tibetana* and *Populus euphratica* also showed that the δ^13^C value of plants increased with the elevation in Tibet (*Chen, 2010; Aini, 2022*). The change of δ^13^C and WUE values of *C. gigantea* may be due to the high altitude of the mountain slope habitat, which is far from the Yarlung Zangbo River. The gradient of the land is relatively steep, making it difficult for *C. gigantea* to obtain water, which may be lead to an improvement in WUE by adjusting stomatal conductance.

### SOURCE OF WATER ABSORPTION BY *C. gigantea*

The contribution rate of potential water source to *C. gigantea* varies with seasons during the study period. Along the Yarlung Zangbo Riverside, *C. gigantea* primarily absorbed river water during dry seasons with little rain, but switched to soil water when there was enough rainfall in August 2023. However, in August 2022, river water remained the main source of absorption due to less rainfall that year. In the terrace and mountain slope habitats, *C. gigantea* mainly absorbs soil water, accounting for about 60%. There was no obvious pattern in the utilization of water from the upper and lower layers of soil by *C. gigantea* across different seasons. An exception occurred in the terrace habitat, where the proportion of groundwater absorption increased and the proportion of soil water decreased in August 2022. In the mountain slope habitat, in November 2023, the proportion of *C. gigantea* absorbing soil water decreased, while the proportion of river water and groundwater absorption increased. There may be two possible reasons for these observations: (1) In terrace habitat, the increased absorption rate of groundwater in August 2022 may be due to the drought of that year, leading to low soil moisture content and water redistribution in the roots of *C. gigantea*. The relatively thick soil layer in terrace habitats compared to slopes and riverbanks may allow deeper root system to absorb groundwater, and redistributing water within the plant. Further research is needed to confirm this. 2) In the mountain slope, *C. gigantea* still primarily absorbed soil water, accounting for more than 60%, which may be related to the high WUE of *C. gigantea* in these the habitats. Precipitation is usually the main source of soil moisture replenishment (*Zhao & Cheng, 2001*). From this point, it can be deducted that the source of water absorption by plants is closely related to their habitat. When the soil moisture content is high, *C. gigantea* will switch to absorbing and utilizing soil water. However, long-term monitoring is needed to determine this relationship definitely, as soil moisture is an important medium connecting vegetation and hydrological processes (*Mahmood, 2011*), and is also a key factor limiting vegetation growth in arid areas (*Noy Meir & Imanuel, 1973*).

## CONCLUSION

Overall, the values of TN_soil_, SOC, NO3-N and NH4-N of soil were significantly higher in mountain slope habitats compared to riverside and terrace, while the pH value of mountain slope soil was significantly lower than that of other two habitats. The total nitrogen content and water content of leaves in riverside habitats were significantly higher than those in terrace and mountain slope habitats, while the water use efficiency of leaves were the highest in mountain slope habitats. This study indicated that the main water source for *C. gigantea* in the distribution area is soil water in terrace and mountain slope habitats, while *C. gigantea* mainly utilized river water at riverside habitats.

## ADDITIONAL INFORMATION AND DECLARATIONS

### Funding

Meizhen Liu was supported by the National Natural Science Foundation of China for Grants 32071604 for this research. The funders had no role in study design, data collection and analysis, decision to publish, or preparation of the manuscript.

### Grant Disclosures

The following grant information was disclosed by the authors:

the National Natural Science Foundation of China (32071604).

### Competing Interest

The authors declare there are no competing interests.

### Author Contributions

- Yueliang Jiang and Meizhen Liu conceived and designed the experiment, performed the experiment, analyzed the data, prepared figures, and tables, written and reviewed drafts of the article, and approved the final draft.
- Li Luan, Youlu Zuo, Shengyun Liu, Qiaoling He analyzed the data, prepared figures, and tables, written and reviewed drafts of the article, and approved the final draft.
- Ruru Wu, Gaoming Jiang performed the experiment, analyzed the data, prepared figures, and tables, written and reviewed drafts of the article, and approved the final draft.

### Data Availability

The following information was supplied regarding data availability: The data is available on reasonable request.

## Notes

### Competing Interest Statement

The authors have declared no competing interest.

